# Genomic response to sex-separated gene pools

**DOI:** 10.1101/2025.07.18.665591

**Authors:** Chloe Melo-Gavin, Karl Grieshop, Aneil F. Agrawal

## Abstract

Males and females experience differences in the strength and direction of selection but discerning the type of genes that are targets of sex differences in selection is complicated by their shared genome. We used experimental evolution in *Drosophila melanogaster* to partially separate the gene pools of males and females for 130 generations. In six replicate populations, we forced one pool of genetically variable Chromosome 2s to experience patrilinear inheritance (segregating like a Y-chromosome) and male-limited selection. The alternative pool segregated like an X-chromosome and experienced female-biased selection. This allowed alleles which are differentially selected for between the sexes to diverge between these pools, enabling us to gain insight into the type of genes subject to such selection. We find that genes which diverge between these pools have an elevated intersexual genetic correlation(*r_MF_*) for expression on average, consistent with the idea that high genetic correlations may hinder sex-specific adaptation under normal inheritance. Diverged genes were also enriched for moderately male-biased genes whereas female-biased genes were underrepresented. At the SNP level, we find an overrepresentation of diverged SNPs involved in splicing or occurring in the 5’UTR and an underrepresentation of missense or synonymous SNPs, suggesting sex differences in selection for isoform usage.

## Introduction

Differences in selection between males and females have been noted among many species and across a variety of phenotypes (Long and Rice, 2007; Abbott et al., 2010; Okada et al., 2021; Arnqvist and Rowe, 2005; Hawkes et al., 2016; Bonduriansky and Rowe, 2005). However, the sexes share most of their genome, and the genetic basis of most homologous traits is highly correlated between them (Poissant et al., 2010; Griffin et al., 2013; Allen et al., 2018), which can hinder their ability to adapt to these different selection pressures. The most obvious example of this is intralocus sexual conflict (IASC), where alleles at a shared locus have opposing fitness effects between the sexes. This antagonistic selection can prevent either sex from achieving their preferred genotype, leading to sub-optimal fitness (Parker, 1979). Even when selection is sexually concordant in direction, differences in the strength of selection means that the more strongly selected sex has its response to selection diluted through the shared gene pool, hindering adaptation (Parker, 1979).

Experimental evolution with sex-limited selection has been used several times to infer how the shared genome constrains sex-specific adaptation and the evolution of dimorphism in traits or gene expression (Prasad et al., 2007; Abbott et al., 2010; Abbott et al., 2020; Norden et al., 2023; Rice, 1996; Rice, 1992). Yet it has been underutilized in investigating genomic targets of sex differential selection, including sexual antagonism. Indeed, identifying the type of genes subject to sex differential selection and IASC is an on-going challenge (Mank, 2017). Several methods have been used to attempt to characterize sex differences in selection across the genome from using population genetic metrics to infer sex differential selection (Cheng and Kirkpatrick, 2016; Sylvestre, et al.., 2023; Ruzicka et al., 2022) to testing associations of alleles or expression patterns with sexually antagonistic fitness effects (Innocenti and Morrow, 2010; Ruzicka et al., 2019). Yet, while evidence is mounting that sex differential selection is pervasive, much remains to be learned about which kinds of genes may be subject to sex differential selection.

Here, we performed sex-limited experimental evolution in *Drosophila melanogaster* over 130 generations to characterize genomic regions subject to sex-differential selection. A pool of genetically variable copies of Chromosome 2 were labelled with a fluorescent *DsRed* marker (hereafter “*Red* chromosomes”). These *Red* chromosomes were experimentally sorted to be inherited from father to son, segregating like a Y chromosome and experiencing male-limited selection, whereas the complimentary pool of non-fluorescent chromosomes (hereafter “*NonRed* chromosomes”) were sorted to segregate like an X, spending ∼2/3 of their time in females and experiencing female-biased selection. This sex-limited selection released *Red* chromosomes from the constraint of the shared genome, allowing male-beneficial alleles to increase in frequency regardless of their effect in females and likewise allowing *NonRed* chromosome pool to increase female-beneficial alleles.

Indeed, males possessing a male-selected *Red* chromosome (hereafter “*Red*” males), displayed increased mating success compared to *NonRed* males, suggesting that separating these gene pools allowed males to adapt to male-specific selection for mating success (Grieshop et al., 2025). Moreover, these two pools of chromosomes had substantial transcriptional divergence (Grieshop et al., 2025), indicating sex differential selection pressures that are normally hindered by the shared gene pool. Data on female fitness is lacking, precluding direct evidence that sexual antagonism is a major source of the sex differential selection. Nonetheless, a major role for sexual antagonism seems likely. The experimental populations were founded from a population well adapted to the lab environment (∼90 generations of lab adaptation with marker segregating before the start of the experiment and more than 230 generation of lab adaptation prior; see Grieshop et al., 2025 for full history) so it is expected that most segregating sexually concordant variation should be fixed or rare and segregating at low frequencies. This means divergence (and patterns of divergence) between these pools was likely dominated by sexually antagonistic variants (see Discussion for further comments). Our aim is to characterize the evolutionary response to sex-biased selection at the genomic level, examining whether the likelihood of allelic divergence between chromosome pools (indicative of the relative likelihood of sex differential selection and of sexually antagonistic selection) is related to properties at the site or gene-level. For example, the level of sex-bias in a gene’s expression may affect the likelihood of a gene experiencing ongoing IASC—as sexual dimorphism (such as dimorphism in expression) has the potential to “resolve” IASC. Cheng and Kirkpatrick (2016) hypothesized that while extremely-sex biased genes could indicate sufficient dimorphism to resolve conflict, and unbiased genes may experience mostly sexually concordant selection, genes with intermediate sex-bias should experience the most ongoing constraint from the shared genome. Indeed, they found evidence from flies that genes with intermediate levels of sex-bias were enriched among candidates for sexual antagonism compared to unbiased or extremely biased genes. However not all studies of sexual antagonism find enrichment of intermediately sex-biased genes (Ruzicka et al., 2019, Ming et al., 2025).

In addition, high intersexual genetic correlations, *r_MF_*, for phenotypes or gene expression hinder the sexes from evolving independently (Lande 1980). When sex differences in selection do occur, genes with high *r_MF_*may be prevented from responding in ways that are ideal for both sexes and, consequently, sexual conflict is more likely to persist. Yet, there is little empirical evidence that high intersexual genetic correlations actually hinder sex-specific adaptation. While intersexual genetic correlations tend to be lower for more strongly dimorphic traits (Poissant et al., 2010) including gene expression (Griffin et al., 2013, but see Singh and Agrawal, 2023), the direction of causality is unclear. That is, traits with high *r_MF_*could be unable to evolve dimorphism or, alternatively, *r_MF_* could decline after dimorphism evolves.

We find that moderately male-biased genes are overrepresented in our diverged genes, implying that they are likely to be constrained by the shared genome under normal inheritance. In contrast, moderately female-biased genes are underrepresented. In addition, our results support the idea that high intersexual genetic correlations for gene expression can constrain genes from resolving sexual conflict. Finally, our results indicate that the functional consequence of a SNP impacts its likelihood to be constrained by the shared genome.

## Methods

### Experimental Evolution Protocol

A detailed experimental protocol is given elsewhere (Grieshop et al., 2025). In brief, six replicate populations of *Drosophila melanogaster* (500 males and 500 females per population each generation) were established from a large (N ≍ 2000) genetically variable lab population originating from the outbred Dahomey base population originally collected from modern-day Benin, West Africa, in 1970. These six populations had been segregating for a dominant fluorescent *DsRed* marker on Chromosome 2 for ∼90 generations prior to the start of the experiment. The marker is located at approximately 2R:48C and no fitness effects of the marker were detected in previous work (Zikovitz and Agrawal, 2013 Supplemental Materials). Hereafter, we refer to copies of Chromosome 2 with and without the *DsRed* marker as “*Red*” and “*NonRed*” chromosomes, respectively. The males used to sire each subsequent generation were *Red*/*NonRed* heterozygotes and females were *NonRed* homozygotes. That is, *Red* chromosomes were transmitted father to son where, due to a lack of meiotic recombination in males, they experienced “Y-like” inheritance and male-limited selection. Conversely, *NonRed* chromosomes (spending 2/3 of the time in females) experienced “X-like” inheritance and thereby female-biased selection. A small fraction of each population underwent slightly altered protocols to allow for a low level of recombination on *Red* chromosomes, both among *Red* and between *Red* and *NonRed* chromosomes. (Since *NonRed* homozygote females are used in the main cross each generation, *NonRed* chromosomes recombine with one another at a higher rate). The expected time a sampled allele spends in either males or females or on *Red*/*NonRed* chromosomes can be estimated by simulation. For example, an allele that is a recombination distance *r* = 0.3 from the *DsRed* marker and sampled from a Red chromosome at generation 130 (when samples were collected for sequencing) is expected to have spent 114.9 (15.1) generations on *Red* (*NonRed*) chromosomes, 115.7 (14.3) generations in males (females), and experienced 0.72 recombination exchanges between *Red* chromosomes, 0.84 between *Red* and *NonRed* chromosomes, and 2.8 between *NonRed* chromosomes. In contrast, a homologous allele sampled from a *NonRed* chromosome is expected to have spent 4.7 (125.3) generations on *Red* (*NonRed*) chromosomes, 46.2 (83.8) generations in males (females), and experienced 0.06 recombination exchanges between *Red* chromosomes, 0.41 between *Red* and *NonRed* chromosomes, and 24.8 between *NonRed* chromosomes.

### DNA extraction and Sequencing

At generation 130, DNA was extracted using DNeasy Blood and Tissue Kit spin column (Qiagen). For each of six populations, two genotypes were sampled: homozygoous NonRed/NonRed females and heterozygous *Red*/*NonRed* females; 500 females were pooled per each sample. Whole genome sequencing was done with Illumina NovaSeq 6000 with an average read length of 151 base pairs. Sequence data quality was checked and cleaned using *fastqc* and *trimmomatic* (Andrews, 2010; Bolgher et al., 2014). Reads were aligned to *D. melanogaster* r.6.5.2 with *bowtie2* (Langmead et al., 2018), and filtering was done using a PHRED quality score of 30 or higher with *samtools* v1.18 (Li, 2011). We kept sites with a coverage between the 1^st^ and 98^th^ percentile (i.e., a total of 3% of sites were excluded because they had too high or too low coverage). Allele frequencies were obtained using *popoolation2* (Koffler et al., 2011) and vcf files were created using *bcftools* v1.18 (Danecek et al., 2021). Tri-allelic sites, sites with a minor allele frequency (averaged across all populations) smaller than 0.05, and SNPs that were segregating in less than 4 populations were filtered out.

### Differentiation between Chromosome Pools and the Creation of a Null Distribution

In general, pinpointing the true targets of selection via divergence analysis in evolve-and-resequence studies has proven to be very difficult because many SNPs diverge due to linkage disequilibrium with the (unknown) true targets of selection or because of drift (Nuzdhin and Turner, 2013; Franssen et al., 2015). A variety of methods exist to identify significantly diverged SNPs in such studies, and these vary in their strengths and weaknesses (Schlötterer et al., 2015). Some of the more sophisticated methods attempt to account for drift to better identify true targets of selection, but these methods require experimental designs that do not match ours (Spitzer et al., 2020). Our goal here is simply to identify a set of SNPs that have diverged in frequency between *Red* and *NonRed* samples, recognizing that there can be multiple reasons for this divergence. Rather than focusing on the identity of individual SNPs (many of which will not be true targets of selection), we are interested in features emerging from the candidate list as a whole, because only selection (not drift) is expected to cause the candidate variants to share features compared to a random set of SNPs. Thus, the resulting set of candidates is not expected to contain only true targets of selection as both (statistical) linkage to targets of selection and divergence by drift can contribute to this set of candidates. Nonetheless, the set of candidates should be enriched for true targets of selection relative to randomly chosen SNPs and only selection on targets is likely to cause patterns among the candidates relative to non-candidates.

With this goal in mind, candidate diverged SNPs were identified as follows. Counts of the reference and alternative alleles for each SNP were analyzed using a quasi-binomial general linear model, considering both Sample Type (*NonRed* homozygotes or *Red*/*NonRed* heterozygotes) and Population as fixed effects using the *stats* package in *R*. We considered as candidates SNPs for which the Sample Type term was ‘significant’ after employing a false discovery rate of 5% calculated using the *p.adjust* function in *R*. As a surrogate for evaluating divergence by drift, the data were re-analyzed after reassigning Sample Type labels within a population (i.e., swapping *Red* and *NonRed* labels) for three of the six populations. The same statistical procedure as used to determine true candidates (as described above) was used to identify label-swap generated pseudo-candidates. This produced ∼95% fewer significant SNPs than we observed using the real labels (SUPPLEMENTAL TABLE 1; see also Supplementary Text).

For gene level analyses, candidate diverged SNPs were converted to genes using *bedtools* v 2.30 (Quinlan and Hall, 2010) and any gene containing a candidate SNP was considered a diverged gene. Because linkage causes diverged SNPs to be spatially related across the focal chromosome, we built a null distribution of pseudo-candidate genes accounting for this spatial relationship. All candidate SNPs on chromosome 2 were shifted in position by the same, randomly chosen amount (where positions shifted past the end of the chromosome loop back to the beginning of chromosome 2) and their new position recorded. This was done 1000 times. In each of these 1000 random location shifts, SNPs were then mapped to their corresponding genes (using *bedtools* as above), creating 1000 lists of pseudo-candidate genes, which were used as the basis for creating null distributions. This approach also accounts for the issue that longer genes are more likely to harbour a diverged SNP simply due to their increased size.

### Divergence in relation to Sex-Biased Expression and rmf

Sex-biased gene expression (SBGE) was measured as fold change (log_2_ male/female) between the sexes (“Log_2_FC”) calculated by Singh and Agrawal (2023) using whole-body transcriptomic data from Osada et al. (2017). Several approaches were used to examine diverged genes with respect to SBGE.

A null distribution of SBGE (i.e., a density distribution of log_2_FC) was constructed as follows. For each of the 1000 location shift pseudo-candidate lists described above, density distributions of SBGE were constructed using the *density* function in base R, which calculates density for 512 evenly spaced values across the range of log_2_FC values (spanning from -10.42 to 14.45). For each of these 512 values, the median density value (from the 1000 random location shifts) was used to construct the null density distribution (and the 2.5th and 97.5th percentile values were used to establish 95% bootstrap confidence intervals). This was then compared to the density distribution of log_2_FC values from diverged genes to determine whether the observed density fell outside the confidence intervals from the null.

The magnitude of allele frequency divergence between two groups could be affected by the starting frequency of the ancestor. To confirm that differences in SBGE between null and diverged genes were not sensitive to starting allele frequency, the above density distribution was also done in allele frequency windows, using frequency data from the earliest generation available, generation 7. The generation 7 allele frequencies were calculated for each SNP using *popoolation2* as above with one exception. Because sequencing data for generation 7 has very low coverage per sample, read count data was pooled across both sample types and across all populations; the resulting pooled count was intended as our best available proxy for the global ancestral allele frequencies. Only SNPs with more than 20 counts after pooling were included.

SNPs were assigned to bins with respect to minor allele frequency (bin size 0.1). For each ancestral minor allele frequency (MAF) bin, genes were considered to have diverged if the “Sample Type” term (i.e., *Red*, *NonRed*) was significant for at least one SNP in the focal MAF bin. Genes with no SNPs within the focal MAF bin were excluded. Density distributions of SBGE values were compared between diverged and non-diverged genes for each MAF bin.

For some analyses, sex-biased genes were sorted into seven categories by degree of bias: extreme female bias (log_2_FC < -5, strong female bias (-5 <log_2_FC< -2), moderate female bias (-2 < log_2_FC <0.5), unbiased -0.5 < log_2_FC <0.5), moderate male bias (0.5 < log_2_FC < 2), strong male bias (2 < log_2_FC < 5), extreme male bias (log_2_FC > 5). A 2×7 χ^2^ test was conducted to test for an association between divergence status (i.e., “diverged” or “not”) and SBGE. We then performed a follow-up test to identify which SBGE categories were over/under-represented among diverged genes as follows. For each of the 1000 location shift pseudo-candidate lists, the fraction of genes in each SBGE category was calculated. Of the 1000 fractions for each category, the 97.5^th^ and 2.5^th^ quantiles were then used to establish 95% confidence intervals for each SBGE category under the null hypothesis. This was then compared to the observed fraction of diverged genes in each SBGE category.

The candidates were examined with respect to the intersexual genetic correlation for expression (*r_MF_*) using *r_MF_* values taken from Singh and Agrawal (2023), which were estimated from expression data in Huang et al. (2015). A null distribution for mean *r_MF_*was calculated from the mean *r_MF_* values from each of the 1000 location shift pseudo-candidate lists and compared to the observed mean *r_MF_* of diverged genes. (To check if the pattern with respect to *r_MF_* was influenced by early allele frequency, mean *r_MF_* was compared in allele frequency bins (bin size 0.1) using allele frequency data from generation 7, similarly to above.) To evaluate whether the relationship between diverged genes and *r_MF_* was due to the association of *r_MF_* and SBGE, we stratified the data into the 7 SBGE categories described above. We then calculated null distributions for mean *r_MF_* within each category.

### Predicted Biological Function of SNPs Analysis

The functional categories of variant SNPs were predicted with ensembl Variant Effect Predictor (VEP) (McLaren et al., 2010), using default settings. This was performed only on chromosome 2, using the raw vcf data. SNPs were further filtered to provide one VEP “consequence” for each unique SNP and biotype combination, taking the consequence in the uppermost hierarchy. The categories “upstream” and “downstream” were combined, and all splice site categories (such as splice donor 5^th^ base variant or splice region variant) were treated as one category. Categories with less than 100 variants were excluded from analysis. After this, there were the following categories: intergenic, variants 5 kb upstream or downstream of a gene, noncoding RNA exons, the 3’ untranslated regions, 5’ untranslated regions, splice sites, introns, missense mutations, and synonymous mutations. We first tested whether diverged SNPs showed enrichment for intergenic versus genic (i.e., all other gene categories analyzed) sites using a _X_^2^-test comparing the numbers of diverged and non-diverged SNPs in the two categories. Subsequently, we tested for enrichment among the genic sub-categories using only variants that occurred in (or around) protein coding genes; variants that did not occur around protein coding regions including SNPs in or near non-coding genes were excluded. We performed a _X_^2^-test comparing the numbers of diverged and non-diverged SNPs in these categories. In addition, for each category, we determined the fraction of SNPs that were significantly diverged and established 95 percent confidence intervals for this fraction using the *boot* package in *R*.

### Comparison to D. simulans

We tested for SNPs shared between our dataset and those segregating in a sample of 170 whole genome sequences from North American inbred lines of *D. simulans*, a closely related species (diverged ∼1 million years ago). The *D. simulans* data set was created by Signor et al. (2018), filtering for sites with a call rate > 80%, depth > 10, and removing triallelic sites. VCF files were downloaded from https://zenodo.org/record/154261#.XEHMtM_7TUJ. *D. simulans* genome coordinates were converted to *D. melanogaster* r6 using the liftOver tool (Casper et al., 2018). Because the minor allele frequency distributions differ between our experimentally diverged and non-diverged SNPs, sites were subsampled to create similar minor allele frequency distributions. First, all sites were sorted into minor allele frequency bins (each of a width 0.05) based on generation 130 allele frequency averaged across all populations and both sample types. For each bin, either diverged or non-diverged SNPs (whichever type was more numerous) were randomly subsampled down so that there was an equal number of SNPs in both categories. A _X_^2^-test was conducted comparing diverged and non-diverged sites with respect to whether the same SNP was found to be segregating in *D. simulans*.

## Results

In total we found 37,084 SNPs (FDR < 0.05; SUPPLEMENTAL FIGURE 1), corresponding to 2127 genes (out of 8456 on this chromosome), were significantly diverged between the *Red* and *NonRed* pools after 130 generations in our male-limited vs. female-biased selection regime. This set of genes, having diverged in response to a separation of gene pools between the sexes, is expected to be enriched for those genes previously constrained by ongoing intralocus sexual conflict. A GO enrichment analysis revealed these experimentally diverged genes were enriched for a variety of functional categories (*p* < 0.001; see **SUPPLEMENTAL)**, including several affecting traits previously identified as likely targets of sexually antagonistic selection such as development, wing morphology, genital morphology, and locomotion (Long et al., 2012; Prasad et al., 2007; Abbott et al., 2010). There is significant overlap between diverged candidate genes derived here and those derived from transcriptomic data in Grieshop et al. (2025) from these same populations ( ^2^ test; p<0.001; **SUPPLEMENTARY TABLE 1**).

We next considered the diverged genes with respect to sex-biased gene expression (SBGE). Cheng and Kirkpatrick (2016) predicted that genes with intermediate levels of expression bias—either female- or male-biased—are the most likely to experience ongoing intralocus sexual conflict compared to either unbiased or extremely biased genes (but see Ming et al., 2025). As such we compared the distribution of sex-bias among diverged genes to a null distribution. Specifically, if intermediately sex-biased genes are more likely to experience ongoing IASC they should be enriched among diverged genes compared to null. Overall, patterns among diverged genes differed significantly from null, both when examined on a continuous scale (**FIGURE 1**), or when broken up into discrete SBGE categories (**SUPPLEMENTAL FIGURE 1;** ^2^ test; *p* < 0.01). For intermediate (moderate to strong) levels of sex-bias, diverged genes show an enrichment of male-biased genes but a deficit of female-biased genes, consistent with male-biased genes being more likely to experience ongoing sexual conflict, while selection on female-biased genes seems to prevent divergence between gene pools. The proportion of extremely sex-biased genes (|log_2_FC| > 5) shows little difference between the diverged genes and the null, consistent with the idea that sufficiently high levels of SBGE may “resolve” IASC. We also performed the above comparison in bins based on a proxy of ancestral allele frequency (**SUPPLEMENTAL FIGURE 2**). The patterns were qualitatively similar to those observed in **FIGURE 1**.

**FIGURE 1:**
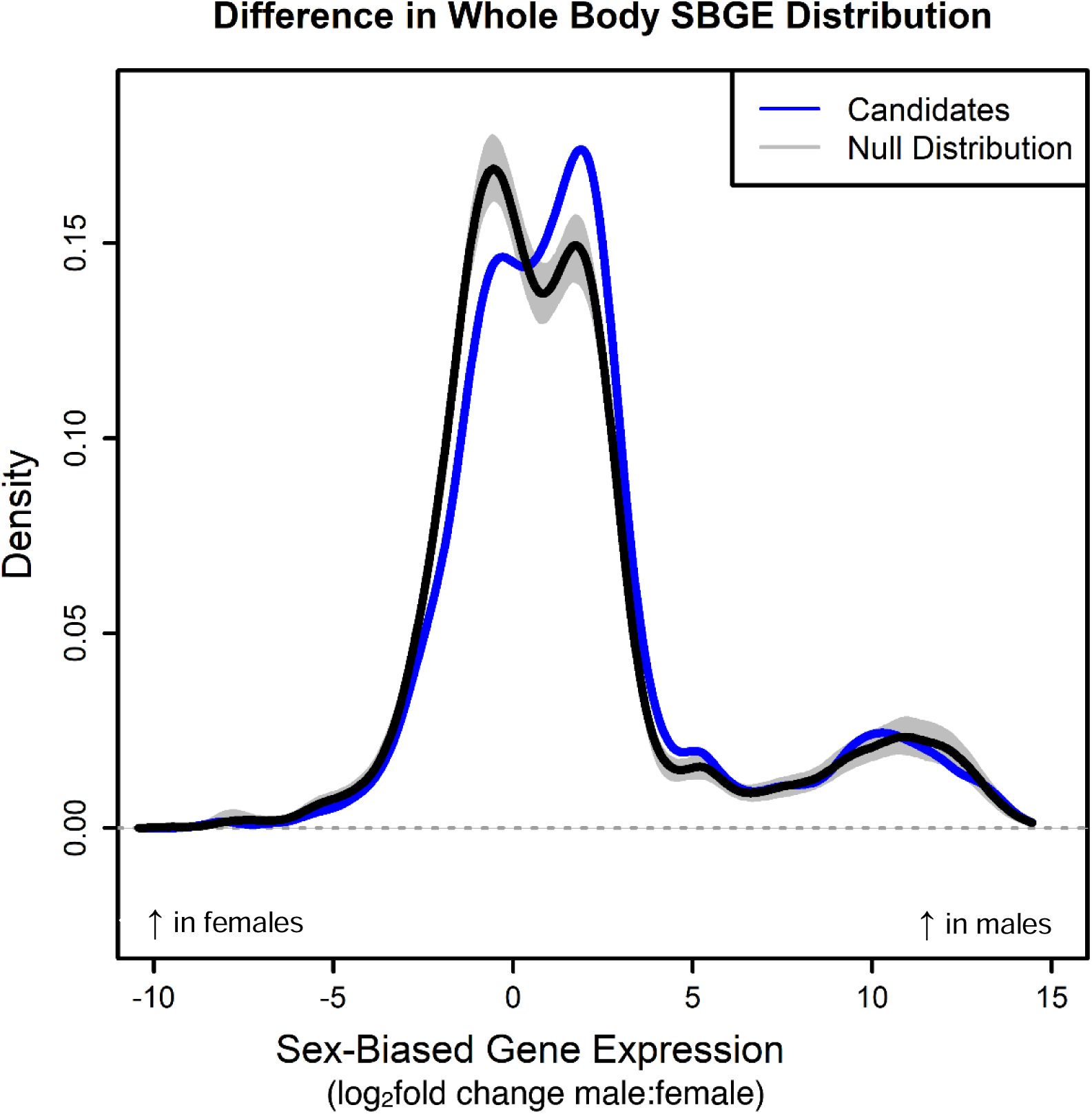
Distribution of Sex-Biased Genes Expression (SBGE) of diverged genes compared to null distribution. Sex-biased gene expression per gene was determined by an external dataset (Singh and Agrawal, 2023) as log_2_FoldChange male:female. The blue line represents the density distribution of diverged candidate genes. The black line represents the median SBGE density distribution of the pseudo-candidates resulting from all 1000 random location shifts (i.e., null distribution), with the grey shaded area representing the 95% confidence intervals. The observed (blue) distribution is considered significantly different than the null given that it lies outside the confidence interval of the null distribution.

Next, we examined diverged genes with respect to the intersexual genetic correlations for expression (*r_MF_*), as measured from an external dataset (Singh and Agrawal, 2023). Because a higher correlation impedes the sexes from evolving independently, we expect genes with stronger intersexual genetic correlations (*r_MF_*) will be more likely to be in a state of unresolved intralocus sexual conflict under normal inheritance and thus more likely to respond to our experimental manipulation of inheritance. Indeed, the mean *r_MF_* of diverged genes is significantly larger than that of the null distribution (*p* < 0.05; **FIGURE 2**). Above we reported that diverged genes are enriched for male-biased genes and a previous study (Singh and Agrawal, 2023) found that male-biased genes have higher *r_MF_* values than unbiased or female biased genes. To evaluate whether the high average *r_MF_* among the diverged genes is due to enrichment of male-biased genes, comparisons of mean *r_MF_* between diverged genes and the null were made separately within seven sex-bias categories (**FIGURE 2**). Diverged genes had significantly elevated *r_MF_*in two main categories: unbiased and moderately female-biased genes (*p* < 0.05), indicating a signal of elevated *r_MF_* independent of the enrichment of male-biased genes. That mean *r_MF_* in diverged genes was elevated in categories with lower sex-bias is consistent with the idea that high *r_MF_* constrains the resolution of IASC.

**FIGURE 2:**
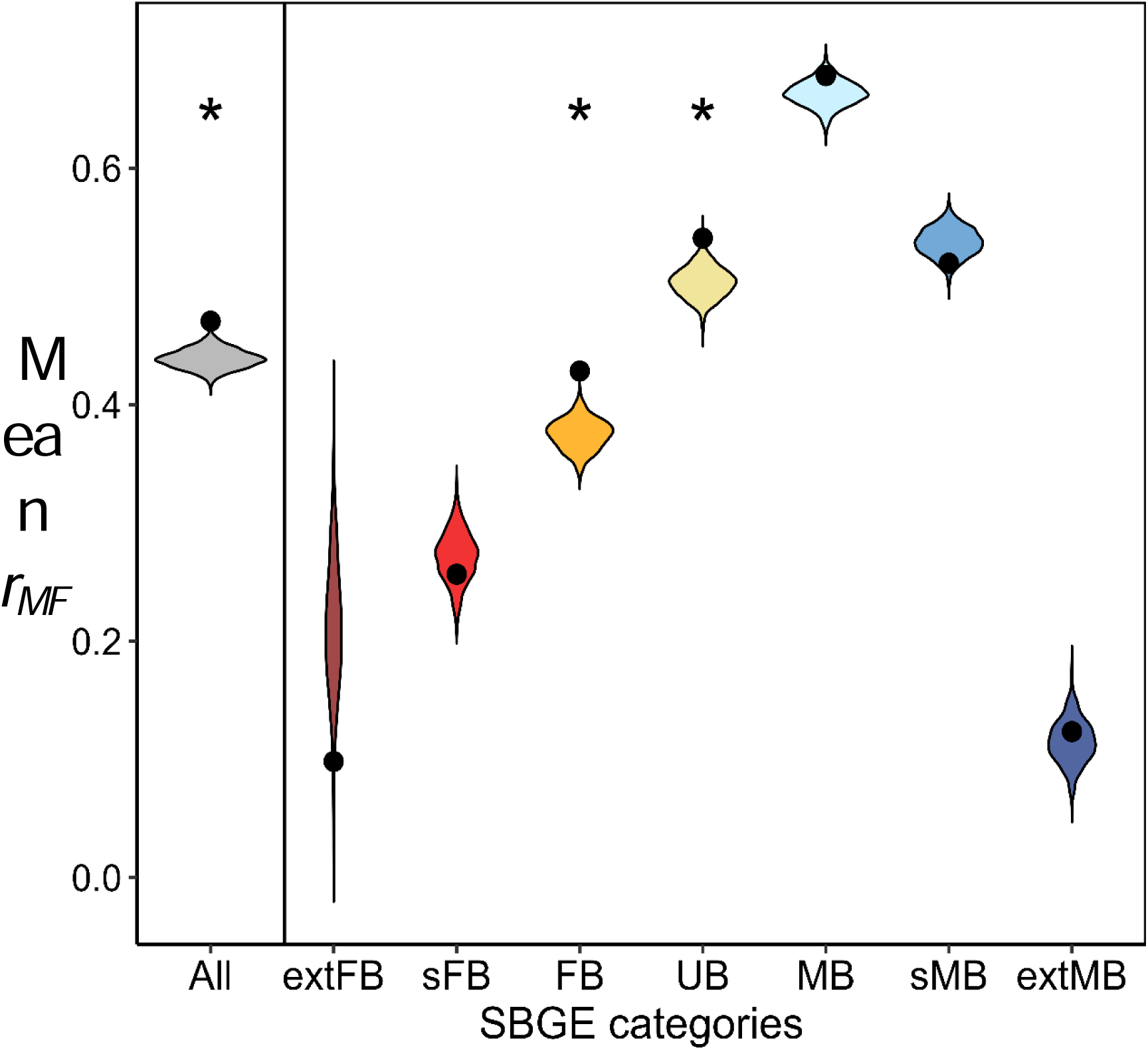
Average intersexual genetic correlation (*r_MF_*) compared between diverged genes and null distribution. Violin plots represent the null distribution of mean *r_MF_* within each category. Black points show mean *r_MF_* of candidate diverged genes within each category. Genes were classified into sex-biased gene categories based on an external dataset. Classifications break down as follows: ALL representing all genes on chromosome 2. Extreme female bias (log2FC <−5), strong female bias (−5 ≤log2FC < −2), moderate female bias (−2 ≤ log2FC < −0.5), unbiased expression (0.5 ≤ log2FC < 0.5), moderate male bias (0.5 ≤ log2FC < 2), strong male bias (2 ≤ log2FC < 5), and extreme male bias (log2FC ≥ 5). Asterisks indicate the mean *r_MF_* for diverged genes is significantly different from the null distribution, lying outside of the 97.5^th^ percentile.

We next examined the biological functions of diverged SNPs. The sexes can experience differences in selection within protein coding sequences as well as regulatory elements but resolving sexual conflict over protein sequence may be more difficult than diverging in expression, resulting in a higher likelihood of experiencing ongoing conflict under normal inheritance. Indeed, a previous study found an excess of protein coding variants, and not regulatory variants, among candidates for sexually antagonistic selection (Ruzicka et al., 2019). We examined our diverged SNPs for enrichment in a variety of functional categories, focusing on variants that occur in intergenic regions, pre-transcriptional regulatory regions (i.e., 5 kb upstream or downstream of a gene), post-transcriptional regulatory regions (e.g., in 5’ or 3’ untranslated region (UTR) or splice sites) or within the protein coding sequence of a gene (including synonymous and missense mutations). Comparing SNPs in intergenic regions to SNPs occurring in genic regions (both protein coding and non-coding), revealed, unexpectedly, that diverged SNPs were significantly enriched for intergenic variants compared to genic variants (*p* < 0.01; **FIGURE 3A**)

**FIGURE 3:**
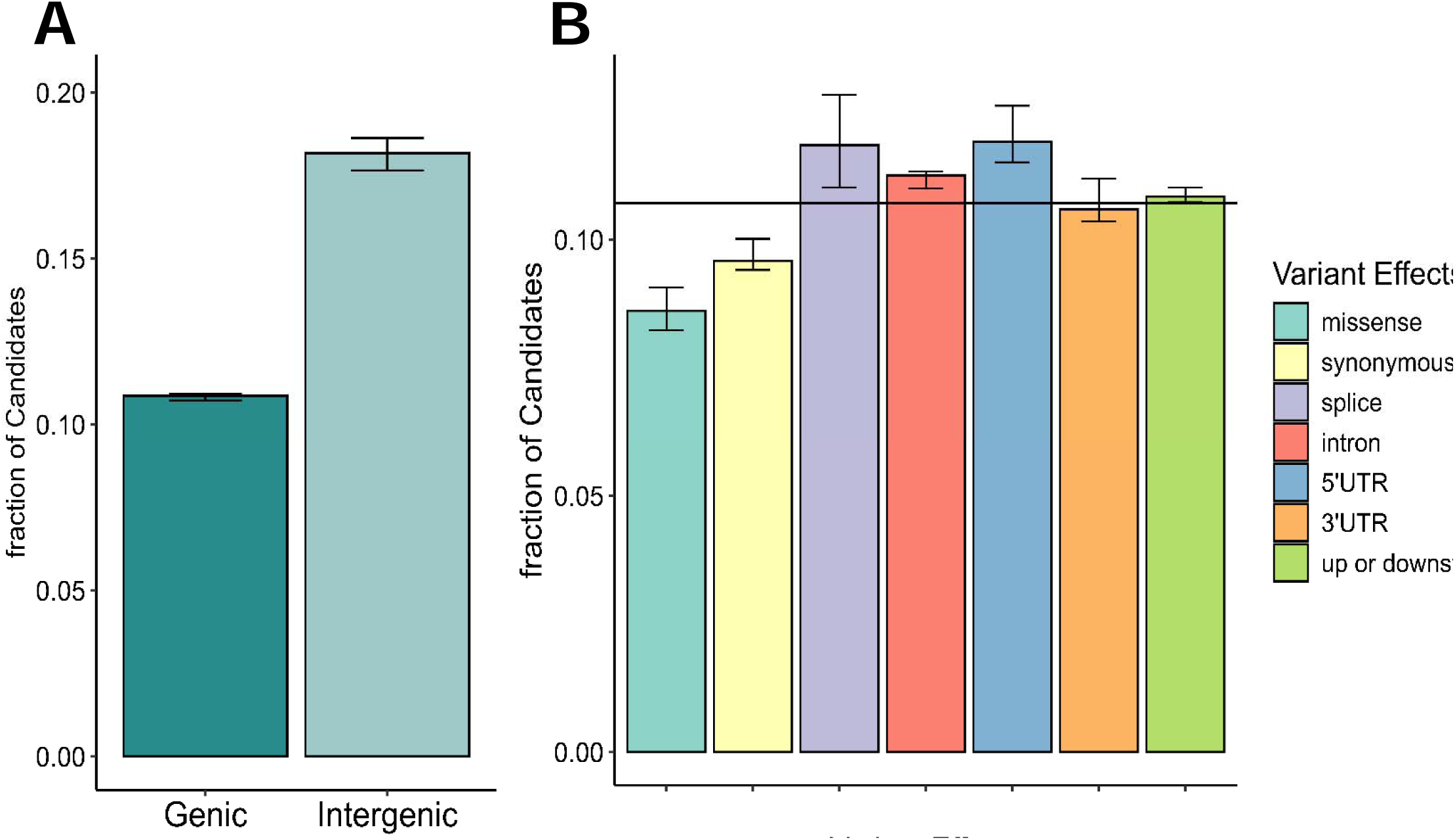
Enrichment with respect to predicted biological function of diverged SNPs. The fraction of candidate SNPs per functional category iscalculated as [number of candidate SNPs in category *i*] / [total SNPs in category *i*]). (A) Comparison of candidate frequency among SNPs occurring in intergenic regions (*n* = 12903) to that among SNPs in or around genes including noncoding genes (genic). (B) Comparison of candidate frequency for each functional category in or near protein coding genes (*n_missense_* = 2670, *n_synonymous_*= 6518, *n_splice_* = 920, *n_intron_* = 47167, *n_5’UTR_* = 1797, *n_3’UTR_* = 3107, *n*_up/downstream=_47661). The solid black line represents unweighted average of the fraction of diverged genes across all 7 categories. Error bars represent bootstrapped 95% percentile confidence intervals.

Considering only variants in or around protein coding genes, we do not see any enrichment for variants occurring within protein coding regions (**FIGURE 3B**). In fact, both missense and synonymous variants are underrepresented among diverged SNPs ( *^2^ test*; p<0.01), contrary to the finding of a GWAS-based analysis of sexually antagonistic variants in flies (Ruzicka et al., 2019). Instead, both splice sites and 5’UTR variants are comparatively enriched for diverged SNPs.

Because IASC has the potential to maintain antagonistic variation via balancing selection, one would predict diverged SNPs might persist over longer evolutionary timescales than background SNPs, possibly spanning speciation events (Ruzicka et al., 2019). However, when assessing whether the divergent SNPs identified in our experiments are also present as polymorphisms in closely related species *Drosophila simulans* (diverged 1 million years ago), this prediction did not hold. In fact, the experimentally divergent SNPs were significantly *less* likely to be found as *D. simulans* polymorphisms compared to background SNPs (chi-squared; x<0.01; S**UPPLEMENTARY TABLE 2**). This suggests that the experimentally divergent SNPs are not targets of long-term balancing selection.

An alternative to sex differences in selection driving divergence between *Red* and *NonRed* is genetic drift. As the experimental design allowed for a small amount of recombination between *Red* and *NonRed* chromosome pools during evolution (see Methods), this would have limited, though not eliminated, the scope for divergence by drift. As a surrogate for the effects of drift, we tested for divergence in permuted data sets in which we reversed the “*Red*” and *“NonRed*” labels in half of the populations. There was an order of magnitude fewer diverged SNPs (‘pseudo-candidates’) even in the most extreme permutations (Supplemental Table 1). When we reduce our observed ‘true’ candidates by the number of pseudo-candidates, the reported patterns remain qualitatively unchanged (see Supplementary Text), providing no support that drift is driving these patterns.

Lastly, the change in allele frequency of diverged SNPs between our proxy for the ancestor (generation 7 sequence pooled across populations and sample types) and generation 130 was compared in both *Red* and *NonRed* sample types. Diverged SNPs changed more over time in the *NonRed* (female-biased selection) samples then in the *Red* (male-limited selection) samples (average magnitude of allele frequency change on Chromosome 2: 8.28 ± 0.03% in *Red* vs 8.63% ± 0.04% in *NonRed*). Thus, there was more evolution within the *NonRed* pool of chromosomes than within the pool of *Red* chromosomes. This could suggest that females are more constrained by a shared gene pool than males or this could be due to *Red* chromosomes suffering more from Hill-Roberston effects due to their lower rate of recombination.

## Discussion

The shared genome can constrain the sexes from adapting to sex-specific selection (Prasad et al., 2007; Abbott et al., 2010). Here, we characterized genes that diverged in response to 130 generations of sex-biased experimental evolution, reasoning that these diverged genes should be enriched for those that experience sex differences in selection, particularly those constrained—under normal inheritance—from adapting to sexually antagonistic selection. Because linkage disequilibrium can be an issue with evolve-and-resequence experiments (Schlötterer et al., 2015), we made no attempt to identify individual targets of selection. Instead, we looked for broadscale patterns of enrichment, which linkage disequilibrium is not expected to produce. Rather, such patterns of enrichment are expected to be due to deterministic forces such as sex differences in selection. While sex differences in selection can occur in both direction (i.e., sexual antagonism) and magnitude (even when direction of selection is concordant), we suspect that sexually antagonistic variants are particularly important. Because the ancestral population used to found the experimental populations had a long prior history in the lab, beneficial sexually concordant variants would likely have been fixed or segregating at high frequencies prior to the start of the experiment and so were unlikely to be major contributors to allele frequency changes between *Red* and *NonRed* pools. Previous work on these populations (Grieshop et al., 2025) showed both mating and transcriptional differences between *Red* (male-selected) and *NonRed* (female-selected) genotypes, suggesting distinct selection pressures between these two chromosome pools. More specifically, they observed increased mating success for male-selected *Red* males, indicating selection for traits affecting this major fitness component that were presumably constrained when gene pools were shared between the sexes. While lacking direct fitness assays on female fitness, Grieshop et al., (2025) also found evidence that male-selected *Red* females have different and plausibly maladaptive mating patterns relative to female-selected *NonRed* females. In addition, here we reported GO enrichment for traits known to be under sexual antagonistic selection (see SUPPLEMENTAL). In sum, we expect, based on the experimental design, that the diverged genes should be enriched for genes subject to sexual antagonism and, though we lack definitive evidence of antagonistic fitness effects, the existing data are consistent with such effects. While we cannot know how strong this enrichment is, our diverged candidate SNPs can be considered candidates for IASC in the sense that they are more likely to experience IASC than background SNPs.

The genomic data provide the opportunity to gain further insights into the transcriptomic divergence previously reported (Grieshop et al., 2025). We find significant overlap between genes which diverged in allele frequency and those which diverged in expression as reported in Grieshop et al., (2025) ( *^2^*-test; *p* < 0.01). This suggests that observed changes in gene expression are likely to be caused by a considerable amount of *cis* regulatory changes. If a large fraction of expression changes were due to a few *trans*-acting factors, then no enrichment would be detectable. The observed overlap between genomic and transcriptomic divergence within Chromosome 2 corroborates the claim of widespread *cis* effects made by Grieshop et al. (2025) based on excess transcriptomic divergence on Chromosome 2 (affected by *cis* and *trans* effects) versus Chromosome 3 (affected by *trans* effects only). A second notable similarity between the genomic and transcriptomic results is that *NonRed* pools show greater changes over time in both allele frequency and gene expression from compared to *Red* pools (Grieshop et al., 2025). This could be due to differences in effective population size, recombination rates, or other differences between *Red* and *NonRed* pools caused by the asymmetrical design of this experiment, or because the sexes may be differentially constrained; that is, they may not be equal distance from their respective optima (i.e., perhaps females are held further from their optimum under normal inheritance and so the female-selected *NonRed* pool shows a larger response to a separate of gene pools).

We find that, for moderately sex-biased genes, both genes that diverged in allele frequency here and those that diverged in expression in Grieshop et al., (2025) were enriched for male-biased genes and had a deficit of female-biased genes. One interpretation of this result is that male-biased genes are more likely to experience ongoing sexual conflict. In the collared fly catcher, Dutoit et al., (2018) found that male-biased genes (but not female-biased genes) had significantly higher intersexual *F_ST_* compared to unbiased genes, potentially indicative of elevated levels of sexual antagonism. Previous work in flies (Cheng and Kirkpatrick, 2016; Innocenti and Morrow, 2010) found enrichment for both types of sex-biased genes among sexually antagonistic candidates, but the enrichment for male-biased genes was of much larger magnitude. Taken together, this suggests that, relative to unbiased genes, male-biased genes are more likely to be constrained from responding to sex-specific selection under normal inheritance.

However, it is possible that moderately female-biased genes experience comparable amounts of sexually antagonistic selection to male-biased genes but do not respond to a separation of gene pools because they experience additional constraints, perhaps due to within-individual pleiotropy. Indeed, female-biased genes are expressed more broadly over spatial and temporal scales than male-biased genes, which are also more tissue specific (most male-biased genes are expressed in the testis) and less networked compared to female-biased genes (Zhang et al., 2007; Allen et al., 2018; Assis et al., 2012; Hansen and Kulathinal, 2013). This could mean that sex-biased experimental evolution disproportionately allows male-biased genes to resolve IASC while female-biased genes remain constrained.

When the intersexual genetic correlation of expression (*r_MF_*) is high, it is harder for the sexes to evolve independently. However, empirical evidence that high *r_MF_* acts to constrain the resolution of sexual antagonism is sparse (Poissant et al., 2010; Griffin et al., 2013; Steward and Rice, 2018). Here, we found that candidate diverged genes had elevated *r_MF_*, suggesting that genes with high *r_MF_* are more likely to experience ongoing conflict under normal inheritance, and that *r_MF_*can act as a barrier to prevent the resolution of IASC.

Notably, male-biased genes, on average, have a higher *r_MF_*than unbiased genes, while female-biased genes have a lower *r_MF_* (Singh and Agrawal, 2023; see also Allen et al., 2017). This may contribute to differences among different categories of sex-biased genes in the potential to resolve sexually antagonistic selection on expression under normal inheritance. While the average *r_MF_* of diverged male-biased genes did not significantly differ from null expectation for male-biased genes here, it is important remember the null expectation for male-biased genes is higher than the genome-wide average. Grieshop et al., (2025)’s expression analysis on these same populations found significantly elevated *r_MF_* for their male-biased candidate genes compared to male-biased background genes. Within unbiased and moderately female-biased genes, those which did diverge in our experiment had significantly elevated *r_MF_* compared to their corresponding null expectations. Taken together, these results suggest that the shared genome may pose a greater constraint on male- than female-biased genes, potentially due the former having a higher *r_MF_* on average. Conversely, the deficit of female-biased genes among diverged candidates may be influenced their lower average *r_MF_*, enabling an easier resolution of conflict even under standard inheritance.

Diverged SNPs were non-random with respect to biological function, suggesting different parts of a gene may experience different levels of constraint from the shared genome. Missense variants were underrepresented among diverged SNPs. In contrast, diverged SNPs were enriched for splice site and 5’UTR variants. This could suggest that post-transcriptional regulation might often be an arena for sexual conflict as both of these site classes affect this route. Unlike the 3’UTR which regulates gene expression through processes such as mRNA stability or protein localization (Barret et al., 2012), the 5’UTR primarily regulates translation initiation. Open reading frames within the 5’UTR affect the efficiency of translation and therefore regulate the relative amount of different protein isoforms. In addition, sequences within the 5’UTR can also be translated into the final protein, such as in the case of *sex-lethal* (*Sxl*) which regulates sex determination and dosage compensation in *Drosophila* (Penalva and Sanchez 2003). Combined with the enrichment of splice sites, this may suggest unresolved sexual conflict in protein isoform ratios or exon usage, in addition to the conflict in expression noted above. In line with this, there are splicing pattern differences between the chromosome pools, with *Red* males possessing a more masculinized splicing pattern than *NonRed* males (Grieshop et al., 2025). A study with this species found an enrichment of missense mutations among sexually antagonistic candidate SNPs, and no enrichment in regulatory variants (Ruzicka et al., 2019). It is not clear why our results differ from theirs. One possibility is that the candidates in that study were identified from a GWAS conducted on adult reproductive fitness. As such, it would not capture IASC that involve effects at the larval stage over viability. Though we are aware of no evidence for sexually antagonistic effects within the larval stage alone (Chippendale et al., 2001), antagonism on total fitness can arise from within-sex trade-offs between larval and adult fitness components, even if they are sexually concordant at each life-stage (Zajitschek and Connallon, 2018). Such variants would be not be identified as sexually antagonistic from a GWAS on adult fitness. On the other hand, some alleles with sexually antagonistic effects on adult fitness (potentially captured by Ruzicka et al., (2019)), may have strongly concordant effects at the larval stage such that their net effect on total fitness is neutral or too weakly antagonistic to be selected for under our experimental design.

If IASC prevents sex-specific beneficial alleles from fixing, this constraint could lead to the prolonged maintenance of sexually antagonistic variation (Connallon and Clark, 2012). Indeed, other studies have found enrichment for *trans*-specific SNPs in their dataset (Ruzicka et al., 2019). In contrast, our diverged SNPs were significantly *less* likely to also be found in *Drosophila simulans.* In our case, it is possible that older sexually antagonistic variants are subject to weak balancing selection and may have been lost through processes such as drift in our lab population (Connallon and Clark, 2012). However, our finding of an underrepresentation of conserved candidate SNPs compared to noncandidate SNPs could suggest that *trans*-specific polymorphisms are prevented from diverging between *Red* and *NonRed* pools, perhaps due to another form of balancing selection that is independent of sex (e.g., antagonistic pleiotropy between life stages).

In general, we show that separating the genome using sex-biased experimental evolution results in substantial divergence between chromosomal pools, indicating that sharing a genome can prevent the sexes from responding to divergent selection pressures. In addition, we characterize the types of genes most likely to be constrained by this shared gene pool. We show that sex-biased genes do not respond symmetrically to this evolutionary manipulation, indicating that male-biased and female-biased genes experience different levels of constraint by the shared genome. We also provide empirical evidence that genes with stronger positive *r_MF_* are likely more constrained under normal inheritance. Additionally, we implicate sex differences in selection in splicing and protein isoform usage as being a site of ongoing sex conflict, which has been relatively understudied until this point.

## Supporting information

GO enrichment

## Author Contributions

C.M.G conducted bioinformatic and statistical analyses, wrote and prepared manuscript, preformed fly population maintenance and DNA extraction. K.G extensive maintenance of fly population and DNA extraction. A.A designed experimental evolution, data analyses and wrote and prepared manuscript.

## Conflict of Interest

There are no conflicts of interest to report.

## Funding

Funding was provided by National Science and Engineering Research Council of Canada (NSERC) (AFA).

## Acknowledgements

We are grateful for everyone who helped us maintain these populations.

## Data availability

DNA-Seq reads will be deposited in the Sequence Read Archive (SRA) upon acceptance. All code and data for analyses and figures are documented in the following GitHub repository: https://github.com/cmmelog/Sex-Biased-Experimental-Evo and will be made public upon acceptance.

## Notes

### Competing Interest Statement

The authors have declared no competing interest.

### Summary of Updates

Methods updated and expanded in Supplemental Material

