## Supplemental Materials for "Genomic response to sex-separated gene pools"

SUPPLEMENTAL FIGURES

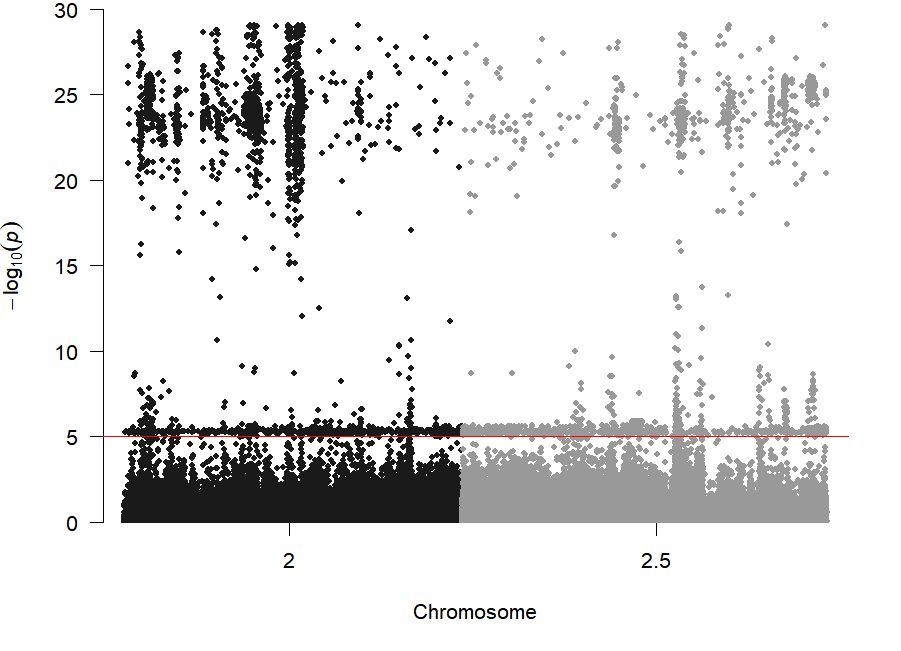

2R

2L

**SUPPLEMENTAL FIGURE 1:** Manhattan plot of SNP association with divergence in allele frequency between *Red* and *NonRed* chromosome pools. (That is, the adjusted p-value for Sample Type (*Red/NonRed)* term in the general linear model) for each variant detected on Chromosome 2. The *DsRed* marker used to mark *Red* flies occurs around 10.7-11.875 of 2R. The blue lines represents a false discovery rate of 5% and the red line represents a false discovery rate of 7.5%.

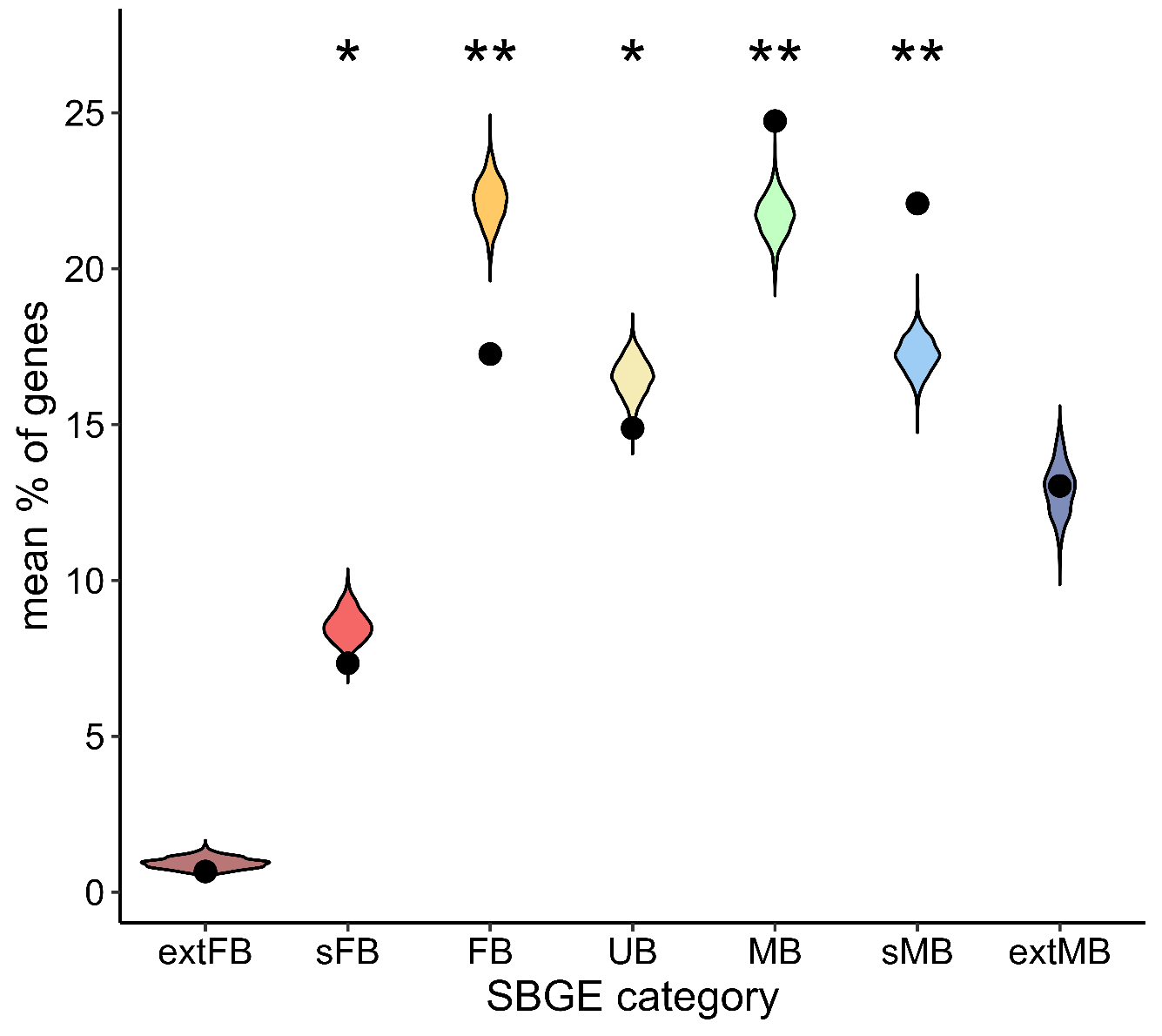

**SUPPLEMENTAL FIGURE 2**: Distribution of sex-biased genes of diverged candidates compared to null distribution. Violin plots represent the null distribution within each sex biased category, and black points represent the bootstrapped mean percent of candidates within each sex-biased category. Genes were classified into sex-biased categories based on an external dataset. Extreme female bias (log2FC male:female <−5), strong female bias (−5 ≤log2FC < −2), moderate female bias (−2 ≤ log2FC < −0.5), unbiased expression (0.5 ≤ log2FC < 0.5), moderate male bias (0.5 ≤ log2FC < 2), strong male bias (2 ≤ log2FC < 5), and extreme male bias (log2FC ≥ 5). Asterisks indicate the mean is significantly different from the null distribution, lying outside of the 95^th^ percent (single asterisk) or 99^th^ (double asterisks) bootstrapped confidence interval of the null.

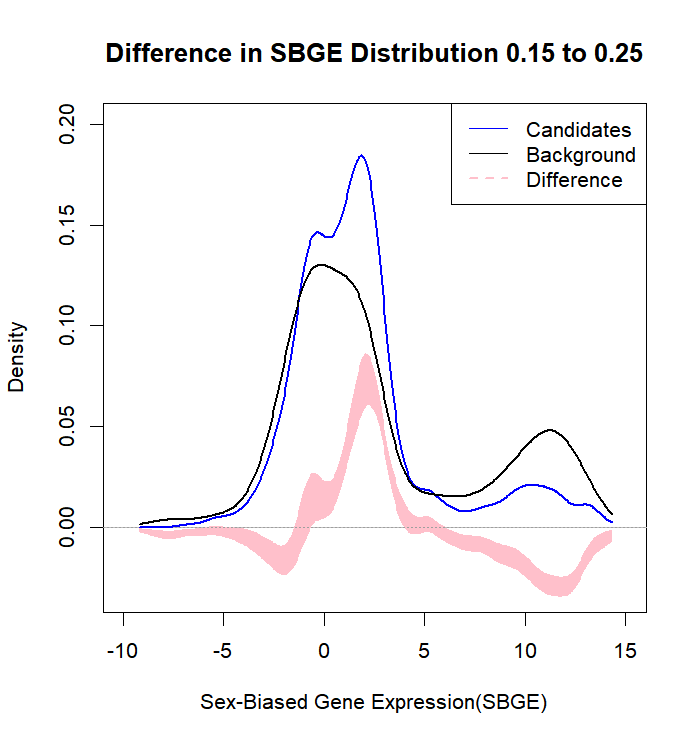

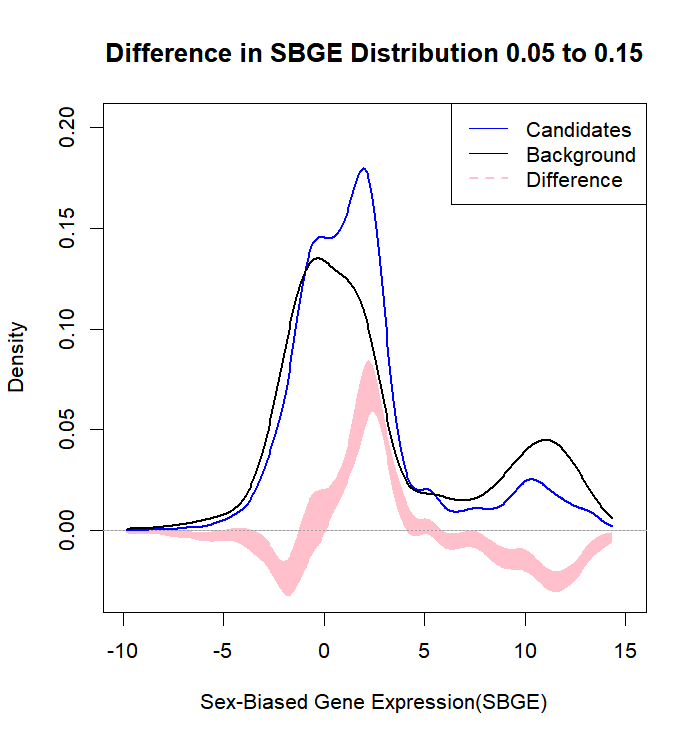

(B)

(A)

(C)

(D)

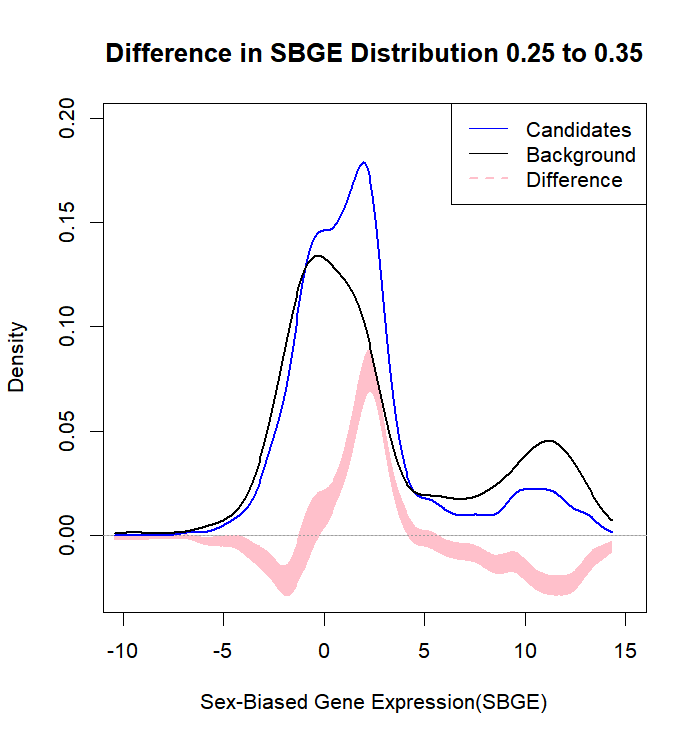

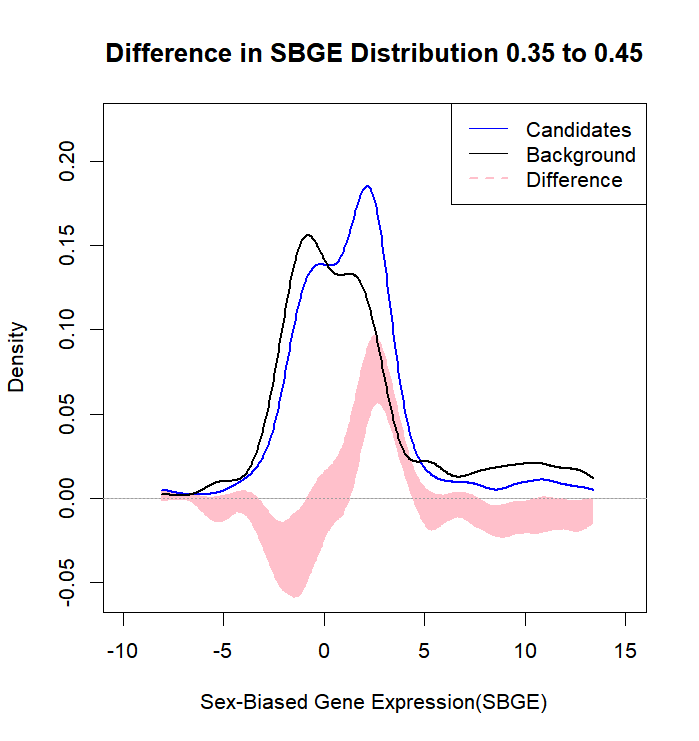

**SUPPLEMENTAL FIGURE 3**: Distribution of sex-biased gene expression (SBGE) of candidate diverged genes compared to background across minor allele frequency (MAF) bins. SNPs were binned based on MAF (bin size=0.1) at Generation 130 and then converted into genes. Sex-biased gene expression per gene was determined by an external dataset (Singh and Agrawal, 2023) as log_2_FoldChange male:female. The blue line represents distribution of diverged candidates genes and black line represents the distribution of background genes. Pink shaded area represents the 95% confidence interval of the difference between the distributions. (A) compares SNPs with MAF between 0.05 to 0.015 (N=1870) (B) compares SNPs with MAF between 0.15 to 0.25(N=1806) (C) compares SNPs with MAF between 0.25 to 0.35 (N=1722) and (D) compares SNPs with MAF between 0.35 to 0.45 (N=520). The observed candidate (blue) distribution is considered significantly different than the background (black) where the 95^th^ CI of their difference (pink shaded area) does not overlap zero.

**B**

**A**

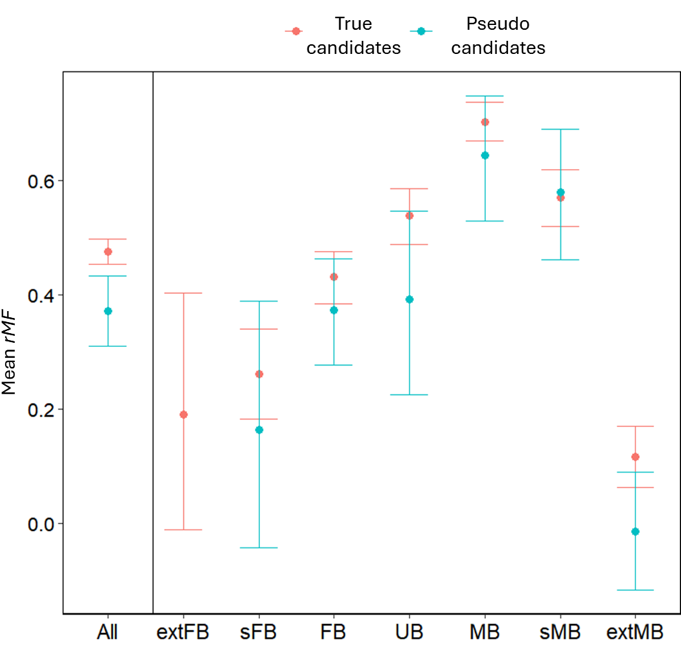

Mean *r_MF_*

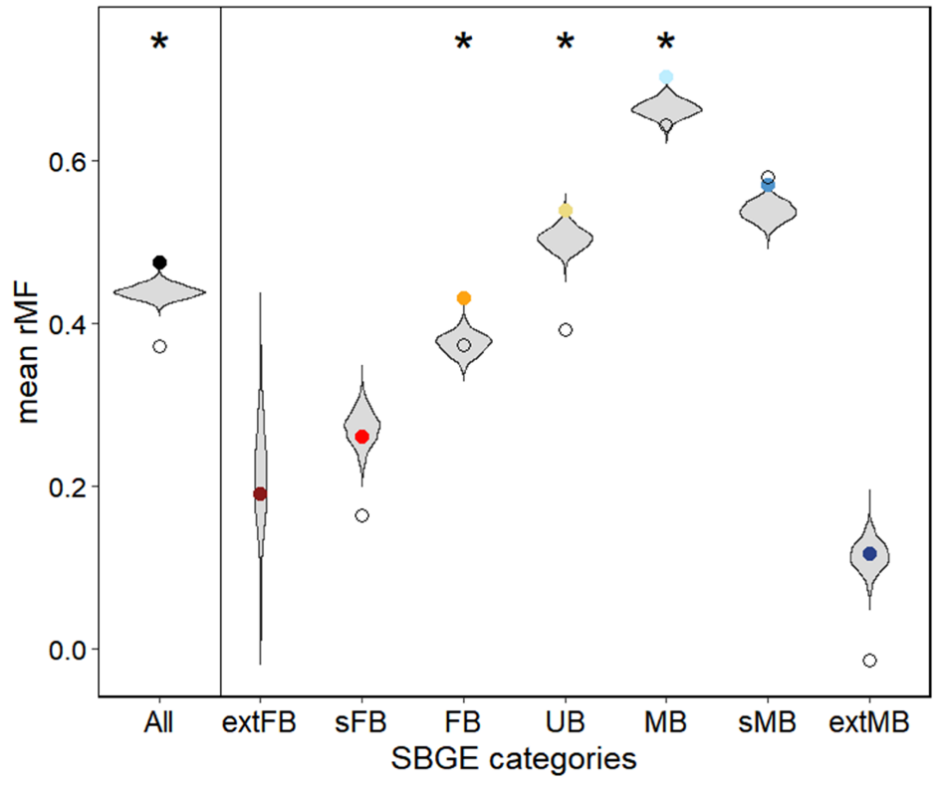

Mean *r_MF_*

SBGE categories

**SUPPLEMENTAL FIGURE 4** Average intersexual genetic correlation (*r_MF_*).. (A) Violin plots represent the null distribution of mean *r_MF_* within each category. Solid circles represent the mean *r_MF_* for unique true candidate genes (i.e., candidate genes after excluding any genes that also appear among the full set of LSG pseudo-candidates). Open circles represent mean *r_MF_* for the unique LSG pseudo-candidates (LSG pseudo-candidates after excluding any genes that overlap with the true candidates). Asterisks indicate the mean *r_MF_* for unique true candidates lies outside of the 97.5^th^ percentile of the null distribution. (B) Mean *r_MF_* between unique true candidates (red) and LSG pseudo-candidates. Error bars represent 95 % bootstrap confidence intervals. Error bars on pseudo-candidates tend to be large because of the low number of pseudo-candidates within most of the sex-bias categories (many fewer than true candidates). Genes were classified into sex-biased gene categories based on an external dataset. Classifications break down as follows: ALL representing all genes on chromosome 2. Extreme female bias (log2FC <−5), strong female bias (−5 ≤log2FC < −2), moderate female bias (−2 ≤ log2FC < −0.5), unbiased expression (0.5 ≤ log2FC < 0.5), moderate male bias (0.5 ≤ log2FC < 2), strong male bias (2 ≤ log2FC < 5), and extreme male bias (log2FC ≥ 5).

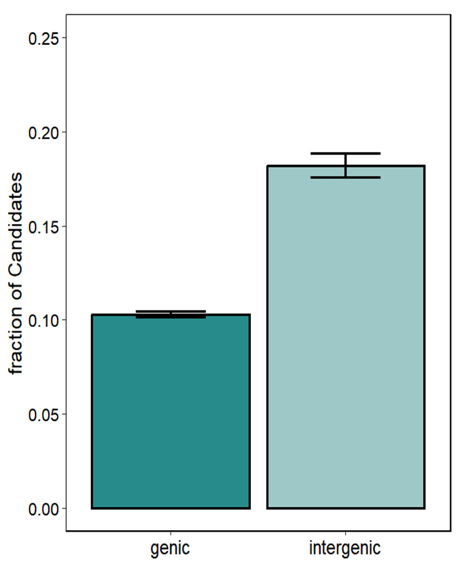

**B**

**A**

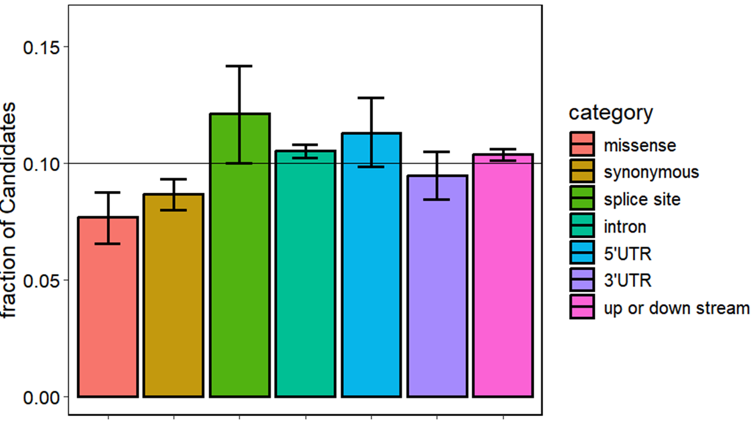

**SUPPLEMENTAL FIGURE 5** Enrichment with respect to predicted biological function of unique true candidate SNPs (candidate SNPs after excluding any SNPs that also appear among the full set of LSG pseudo-candidate SNPs). The fraction of candidate SNPs per functional category (calculated as [the number of candidate SNPs in category *i*]*/* [the total number of SNPs in category *i*]) was calculated per category. (A) Comparison of SNPs occurring in intergenic regions to SNPs with in or around genes, including noncoding genes (genic). (B)Comparison of the fraction of unique true candidate SNPs in each functional category for SNPs in genic regions.

Supplemental Tables

**SUPPLEMENTAL TABLE 1.** Number of Label-Swap Generated (LSG) pseudo-candidate SNPs detected when swapping Sample Type (*Red* or *NonRed*) labels for three of the six populations. Pseudo-candidates were identified using the same statistical criteria as for identifying the true (i.e., non-permuted data) candidate SNPs. Corresponding genes were calculated for each pseudo-candidate SNP.

| Permutation # (swapped populations/  un-swapped populations) | Pseudo-candidate SNPs  (% Overlap* between True Candidates and Pseudo-Candidates) | Pseudo-Candidate genes  (% Overlap* between True Candidates and Pseudo-Candidates) |
| --- | --- | --- |
| Perm 1 (123/456) | 51 SNPs (9.8%) | 29 genes (30.0%) |
| Perm 2 (124/356) | 147 SNPs (6.1%) | 49 genes (52.9%) |
| Perm 3 (125/346) | 697 SNPs (18.9%) | 237 genes (54.4%) |
| Perm 4 (126/345) | 534 SNPs (13.3%) | 233 genes (58.8%) |
| Perm 5 (134/256) | 1310 SNPs (10.4%) | 353 gene (49.3%) |
| Perm 6 (135/246) | 9 SNPs (11.1%) | 8 gene (75%) |
| Perm 7 (136/245) | 4 SNPs (0%) | 4 gene (75%) |
| Perm 8 (145/236) | 13 SNPs (7.7%) | 11 gene (63.6%) |
| Perm 9 (146/235) | 0 SNPs | 0 genes |
| Perm 10 (156/234) | 558 SNPs (15.0%) | 207 gene (58.9%) |

*% Overlap = (number of pseudo-candidates that are also true candidates) / (total number pseudo-candidates)

**SUPPLEMENTAL TABLE 2.** Overlap between diverged candidates identified in this genomic study and those identified via differential expressionin Grieshop et al. (2025). Enrichment determined as presence/absence of our diverged candidate genes within Differentially Expressed (DE) candidate genes of Grieshop et al., (2025). *χ*^2^ test; *χ*^2^=31.81, df=1, p<0.00001

| Overlap with Grieshop et al., (2025) | | | |
| --- | --- | --- | --- |
|  | Background | DE genes |  |
| Background | 3263 | 167 | 4.86% (167/3430)  DE-Genes |
| Diverged Candidates | 1605 | 157 | 8.91% (157/1762) DE-genes |
|  | 32.97% (1605/4868) Diverged Candidates | 51.45% (157/324) Diverged Candidates |  |

**SUPPLEMENTAL TABLE 3.** Comparison of *trans*-specific SNPs occurring in *Drosophila simulans* between our candidate and background SNPs. SNPs within our dataset were compared to 170 North American *D. simulans* (from Signor et al., 2018) to determine *trans*-specific status*.* Candidate SNPs and background SNPs were allele frequency matched before comparison. This was done by downsampling background SNPs so there was an equal number of diverged and background SNPs in each MAF bin (bin size=0.02). *χ^2^*=5.4, df=1, p<0.05.

|  | *Trans*-specific | Not *trans*-specific |
| --- | --- | --- |
| Diverged SNP | 3.07% (1310/42676) | 96.97% (41366/42676) |
| Background SNP | 3.35% (1426/42534) | 96.65% (41108/42534) |

**SUPPLEMENTAL TABLE 4.** Distribution of sex-biased genes. of unique true SNPs.. Genes were classified into sex-biased categories based on an external dataset. Extreme female bias (log2FC male:female <−5), strong female bias (−5 ≤log2FC < −2), moderate female bias (−2 ≤ log2FC < −0.5), unbiased expression (0.5 ≤ log2FC < 0.5), moderate male bias (0.5 ≤ log2FC < 2), strong male bias (2 ≤ log2FC < 5), and extreme male bias (log2FC ≥ 5). Asterisks denotes significant difference from percentage from background based on bootstrapping the difference in percentage

|  | Background | Unique True  Candidates^1^ | Unique  Pseudo-Candidates^1^ | Net  Candidates^2^ |
| --- | --- | --- | --- | --- |
| extFB | 37 (1%) | 8 (0.7%) | 0 | 8 (0.8%) |
| sFB | 253 (9%) | 91 (7.9%) | 13 | 78 (8.2%) |
| FB | 696 (25%) | 211 (**18.3%***) | 37 | 174 (**18.3%***) |
| UB | 439 (16%) | 172 (14.9%) | 26 | 146 (15.4%) |
| MB | 512 (18%) | 283 (**24.6%***) | 45 | 238 (**25.1%***) |
| sMB | 282 (10%) | 218 (**18.9%***) | 37 | 181 (**19.1%***) |
| extMB | 593 (21%) | 169 (**14.7%***) | 44 | 125 (**13.2%***) |
| Total | 2812 | 1152 | 202 | 950 |

^1^Excluding genes that overlap between true and pseudo-candidates

^2^Net = (number of true candidates) – (number of pseudo-candidates)

**SUPPLEMENTAL TABLE 5.** Comparison of genic and intergenic SNPs. Genic sites are underrepresented, and intergenic sites are overrepresented among unique true candidates (*χ*^2^ = 617.4, df = 1, *p* < 10^-15^) compared to background SNPs. The same result holds using the net candidate counts (*χ*^2^ = 562.1, df = 1, *p* < 10^-15^).

|  | Background^1^ | Unique True  Candidates^2^  (% with respect to background) | Unique Pseudo-Candidates^2^  (% with respect to appropriate background^5^) | Net  Candidates^3^  (% with respect to appropriate background^6^) |
| --- | --- | --- | --- | --- |
| Genic^4^ | 114294 | 13911 (10.8%) | 786 (0.1%) | 13125 (10.4%) |
| Intergenic | 10557 | 2346 (18.2%) | 0 (0%) | 2346 (18.2%) |

^1^Background are the compliment of true candidates

^2^Excluding SNPs that overlap between true and pseudo-candidates

^3^Net = (number of true candidates) – (number of pseudo-candidates)

^4^As specified in the main text "Genic" includes upstream and downstream regions surrounding genes as well as introns

^5^Background here are sites that are not pseudo-candidates (either unique pseudo-candidates or those that pseudo-candidates that overlap with true candidates)

^6^Background here are sites that are not true or pseudo-candidates (or those that overlap between true and pseudo-candidates)

**SUPPLEMENTAL TABLE 6.** Enrichment of candidates with respect to predicted biological function for SNPs in genic regions. Within genic regions, unique true candidates are not evenly distributed across functional categories (*χ*^2^ = 41.8, df = 6, *p* = 2.0 × 10^-7^). The same result holds even using the Net Candidate counts (*χ*^2^ = 47.6, df = 6, *p* = 1.8 × 10^-8^).

|  | Background^1^ | Unique True  Candidates^2^  (% with respect to background) | Unique Pseudo-Candidates^2^  (% with respect to appropriate background^5^) | Net  Candidates^3^  (% with respect to appropriate background^6^) |
| --- | --- | --- | --- | --- |
| 3' UTR | 2778 | 329 (10.5%) | 35 (1.1%) | 291 (9.6%) |
| 5' UTR | 1583 | 214 (11.9%) | 12 (0.7%) | 202 (11.4%) |
| synonymous | 5893 | 625 (9.6%) | 64 (1.0%) | 560 (8.8%) |
| Missense | 2440 | 230 (8.5%) | 25 (0.9%) | 203 (7.8%) |
| Splice | 847 | 121 (12.5%) | 4 (0.4%) | 117 (12.2%) |
| intron | 42300 | 5408 (11.3%) | 407 (0.9%) | 4976 (10.6%) |
| Up/down-stream | 56061 | 6719 (10.7%) | 220 (0.4%) | 6496 (10.4%) |

^1^Background are the compliment of true candidates

^2^Excluding SNPs that overlap between true and pseudo-candidates

^3^Net = (number of true candidates) – (number of pseudo-candidates)

^4^As specified in the main text "Genic" includes upstream and downstream regions surrounding genes as well as introns

^5^Background here are sites that are not pseudo-candidates (either unique pseudo-candidates or those that pseudo-candidates that overlap with true candidates)

^6^Background here are sites that are not true or pseudo-candidates (or those that overlap between true and pseudo-candidates)

**SUPPLEMENTAL TABLE 7** Comparison of *trans*-specific SNPs occurring in *Drosophila simulans*. SNPs within our dataset were compared to 170 North American *D. simulans* (from Signor et al., 2018) to determine *trans*-specific status (i.e., whether same polymorphism was present in *D. simulans*)*.* Sites where there is a shared polymorphism with *D. simulans* are underrepresented among the unique true candidates relative to the background (*χ*^2^ = 41.3, df = 1, *p* = 1.3 × 10^-10^). The same result holds with respect to using the net candidates (*χ*^2^ = 44.2, df = 1, *p* = 2.9 × 10^-11^; for this test, the background numbers are appropriately adjusted by counting only those sites that are neither true nor pseudo-candidates).

|  | Background^1^ | Unique True  Candidates^2^ | Unique  Pseudo-Candidates^2^ | Net  Candidates^3^ |
| --- | --- | --- | --- | --- |
| *Trans*-specific | 6639 | 671 | 56 | 615 |
| Not *trans*-specific | 274648 | 36104 | 2348 | 33756 |
| % in *D. simulans* | 2.4% | 1.8% | 2.3% | 1.8% |

^1^Background are the compliment of true candidates

^2^Excluding SNPs that overlap between true and pseudo-candidates

^3^Net = (number of true candidates) – (number of pseudo-candidates)
